# An aromatic substrate prenyltransferase involved in the chemical diversification of flavonoids in *Glycyrrhiza glabra*

**DOI:** 10.64898/2026.05.12.724477

**Authors:** Arisa Kubomura, Takatomo Arai, Junwen Han, Ryosuke Munakata, Noriko Yasuno, Osamu Kobayashi, Kanji Mamiya, Kazuki Nakamura, Nariaki Wasano, Kazufumi Yazaki, Kazuaki Ohara

## Abstract

Prenylated isoflavonoids are widely distributed specialized metabolites within the Fabaceae and contribute to various characteristic biological activities for both plants and humans. Several aromatic prenyltransferases (PTs) have been identified in *Glycyrrhiza* species, which are the most widely consumed crude drugs in traditional Chinese medicine. However, these enzymes do not sufficiently explain the structural diversity of prenylated flavonoids produced in the *Glycyrrhiza* genus. To identify additional novel PTs, we used elicited cultured *Glycyrrhiza glabra* roots as source material, in which elicitor treatment of cultured roots increased the accumulation of multiple prenylated flavonoids. To identify the responsible enzyme, PT candidates were screened using *G. uralensis* transcriptomes, currently the sole publicly available transcriptomic resource within the genus, and a homolog designated GgBSPT1 (BSPT; a broad-substrate prenyltransferase) was subsequently isolated from elicited cultured *G. glabra* roots. GgBSPT1 differed from previously identified *Glycyrrhiza* PTs in both amino acid sequence and enzymatic properties. GgBSPT1 catalyzed 3′-prenylation of isoliquiritigenin and 6-prenylation of five flavonoids, i.e., this PT displayed broad substrate acceptance across 20 distinct flavonoid structures. Overall, elicited cultured *G. glabra* roots enabled the identification of a previously unrecognized PT that is functionally distinct from earlier reported *Glycyrrhiza* PTs. This study provides a new insight into the metabolic plasticity of *Glycyrrhiza* species and expands the enzymatic toolkit for future metabolic engineering of prenylated phytochemicals by the unusually broad substrate specificity of GgBSPT1.

**Main conclusion:** Using cultured *Glycyrrhiza glabra* roots, we identified a new prenyltransferase involved in the formation of a variety of flavonoids, thereby revealing novel prenylated isoflavonoid pathways in licorice.

## Introduction

Plant secondary metabolites comprise a wide spectrum of bioactive compounds with diverse physiological functions. Fabaceae species are particularly recognized for accumulating various isoflavonoids and their prenylated derivatives, which possess considerable structural diversities and display biological activities enhanced by prenyl moieties (Yang and Wang 2024). In licorice (*Glycyrrhiza* species), both triterpenoid saponins, such as glycyrrhizin, and numerous prenylated flavonoids contribute substantially to pharmacological activities, including modulation of glucose and lipid metabolism, cancer prevention, and cardiovascular support (Shao et al. 2025). Among these compounds, glabridin is a prenylated isoflavonoid characteristic of *Glycyrrhiza glabra* that accumulates predominantly in the roots of *G. glabra* (Kondo et al. 2007; Avula et al. 2022). This isoflavone derivative exhibits diverse biological activities, including antioxidant, anti-inflammatory, and anticancer effects, which make it attractive for pharmaceutical, food, and cosmetic applications (Simmler et al. 2013). The biosynthesis of glabridin originates from the general flavonoid pathway and proceeds through tailoring reactions, including prenylation, hydroxylation, and methylation, which collectively generate its characteristic chemical structure and bioactivity, although the precise sequence of these steps remains unclear (Simmler et al. 2013; Lu et al. 2025; Zhang et al. 2026).

Prenylation of aromatic compounds represents a key modification in plant secondary metabolism that generates chemical diversity and enhances biological activity. Most aromatic prenylation reactions are catalyzed by membrane-bound prenyltransferases (PTs) of the UbiA superfamily. These enzymes introduce prenyl side chains into flavonoid scaffolds, thereby modulating hydrophobicity, membrane association, and pharmacological properties (Munakata et al. 2021). Since the first identification of a flavonoid-specific PT in *Sophora flavescens* (Sasaki et al. 2008), approximately 50 UbiA-type aromatic PTs accepting coumarins, phenylpropanoids, and phloroglucinols have been reported from various plant species (Munakata et al. 2021).

Several PTs have been identified in *Glycyrrhiza* species, including GuA6DT (Li et al. 2014), GuILDT (Li et al. 2018), and an GgPT1 (Zhang et al. 2026). However, these enzymes account for only a small fraction of the more than 200 prenylated flavonoids reported in this genus (Li et al. 2014). Therefore, the currently characterized PTs are insufficient to explain the observed structural diversity in licorice. This limited representation suggests that additional, as yet unidentified PTs participate in the biosynthetic pathways of prenylated flavonoids in *Glycyrrhiza* species. Secondary metabolic pathways are differentially activated under distinct cultivation conditions. For example, in vitro plant cultures frequently exhibit altered gene expression and metabolite profiles relative to soil-grown intact plants, reflecting responses to nutrient composition, microbial interactions, and stress signaling (Wawrosch and Zotchev 2021; Fazili et al. 2022; Korenblum et al. 2022; Gupta et al. 2024). The use of in vitro plant cultures or cultured dedifferentiated cells, rather than soil-grown intact plants (Zhang et al. 2026), provides many opportunities to identify novel biosynthetic genes that may remain undetected under conventional conditions. On this basis, we hypothesized that cultured *G. glabra* roots harbor distinct biosynthetic routes of prenylated flavonoid metabolism and may provide access to previously uncharacterized PTs.

In this study, we sought to identify novel PTs in *G. glabra* using cultured roots as the source material. We cloned a new PT gene from elicited cultured licorice roots and characterized its distinctive enzymatic properties, demonstrating its broad substrate and product specificities within a single prenylation enzyme, which suggests the substantial involvement of the new PT in the biosynthesis of various prenylated isoflavones in *G. glabra*. This work uncovers previously hidden branches of specialized metabolism and establishes a foundation for future functional studies, including metabolic engineering of prenylated flavonoids.

## Material and Methods

### Plant materials and cultures

*G. glabra* seeds were kindly provided by the Health Science University of Hokkaido, Japan. Seeds were surface sterilized by immersion in 70 % ethanol for 1 min, followed by treatment with 1 % sodium hypochlorite for 15 min, and subsequently rinsed three times with sterile distilled water. The sterilized seeds were sown on Murashige and Skoog (MS) medium supplemented with 3 % sucrose and 0.2 % gellan gum (pH 5.8). Seedlings were germinated and grown at 25 °C under a 16 h/8 h light–dark photoperiod. After root establishment, plants were transferred to a plant box (SPL Life Sciences, Gyeonggi-do, Korea) containing the same medium and maintained under identical conditions. Cultured plants were subsequently propagated by periodic nodal subculture on MS medium.

### Elicitation of glabridin production in cultured *G. glabra* roots

After the final subculture, roots from twelve 4-week-old cultured *G. glabra* plants were excised, cut into approximately 1-cm segments, pooled, and randomly distributed into plant boxes containing either 50 mL of filter-sterilized 3 % yeast extract solution (Difco Laboratories, NJ, USA) for the elicited group (YE) or 50 mL of sterile water for the negative control group (NC). Cultures were incubated at 25 °C in darkness. Root tissues (50 mg, n = 3) were collected after 1, 3, and 7 days of treatment, designated as days 1, 3, and 7, respectively. Each sample was extracted with 500 µL methanol and homogenized using a Multi-Beads Shocker (Yasui Kikai, Osaka, Japan). After centrifugation at 10,000 × g for 5 min, the supernatant was transferred to a fresh microcentrifuge tube and centrifuged again at 15,000 × g for 5 min. An aliquot (350 µL) of the supernatant was evaporated under reduced pressure and redissolved in 100 µL methanol. The resulting extract was subjected to liquid chromatography-tandem mass spectrometry (LC–MS/MS) analysis.

### Preparation of crude enzymes from cultured *G. glabra* roots

Roots from forty 4-week-old cultured *G. glabra* plants (3 g fresh weight [FW]) were excised, cut into approximately 1-cm segments, and incubated in filter-sterilized 3 % yeast extract solution (Difco Laboratories) for 24 h in darkness. Treated roots were homogenized in 40 mL of 100 mM potassium phosphate buffer (pH 7.0) containing 0.4 g polyvinylpyrrolidone (PVPP) (nacalai tesque, Kyoto, Japan), 10 mM dithiothreitol (DTT) (Fujifilm Wako, Osaka, Japan), and ProteoGuard™ EDTA-Free Protease Inhibitor Cocktail (Roche Diagnostics, Basel, Switzerland). The homogenate was centrifuged at 10,000 × g for 3 min at 4 °C and filtered twice through Miracloth (Merck KGaA, Darmstadt, Germany). The filtrate was subsequently centrifuged at 15,000 × g for 60 min, and the resulting pellet was resuspended in 1 mL of 100 mM Tris-HCl buffer (pH 7.5). After brief sonication, the suspension was diluted tenfold with the same buffer and used as the crude membrane enzyme fraction. Enzyme reactions (total volume 200 µL) contained 0.2 mM flavonoid substrate, 0.2 mM dimethylallyl diphosphate (DMAPP) (Echelon Biosciences, Utah, USA), 10 mM MgCl₂, and 200 µL crude enzyme preparation. Reaction mixtures were incubated for 16 h at 25 °C in darkness. Products were extracted with ethyl acetate, evaporated to dryness, and redissolved in methanol for high-performance liquid chromatography (HPLC) analysis. The HPLC method is described below.

### Identification of candidate PT genes from *G. uralensis*

RNA-seq data derived from underground tissues of *G. uralensis* (SRR18072253, SRR18072254, and SRR18072255) were trimmed using BBduk and assembled using SPAdes within Geneious software version 2023.2.1. Candidate PT contigs were identified by BLAST search using the amino acid sequences of GuA6DT (Li et al. 2014) (Accession No. KJ123716) and GuILDT (Li et al. 2018) (Accession No. KR139751) as queries. From the resulting hits, contigs containing a coding DNA sequence (CDS) longer than 1,000 bp with two conserved aspartate-rich motifs were selected as candidates.

### GgBSPT1 cDNA cloning and construction of expression vectors for *Nicotiana benthamiana*

The CDSs of selected GuPT candidates were synthesized using a custom vector construction service (VectorBuilder, Tokyo, Japan) and cloned under the control of the cauliflower mosaic virus (CaMV) 35S promoter. For cloning of *GgBSPT1* cDNA, total RNA was isolated from cultured *G. glabra* roots (500 mg FW) using the PureLink RNA Mini Kit (Thermo Fisher Scientific, MA, USA). First-strand cDNA was synthesized using the SuperScript IV First-Strand Synthesis System (Thermo Fisher Scientific). Using primers designed from the GuPT1 sequence, GgBSPT1 was amplified by PCR with KOD One PCR Master Mix (Toyobo, Osaka, Japan), employing the forward primer (5′-GGGCATATGGATTCATTGATTGTTGG-3′) and the reverse primer (5′-CCCGTCGACTCATCTAACTAAAGGCATGAG-3′). The amplified fragment was cloned into the NdeI–SalI site of pRI201-AN DNA (Takara Bio, Shiga, Japan) using Ligation high Ver. 2 (Toyobo). The silencing suppressor P19 (accession No. AJ288943) was synthesized using a custom vector construction service (VectorBuilder) and placed under control of the CaMV 35S promoter. The agroinfiltration method for transient expression of foreign genes in *N. benthamiana* is described below.

### Construction of *Saccharomyces cerevisiae* expression vectors

For heterologous expression in *S. cerevisiae*, the transit peptide (TP) region responsible for plastid targeting of GgBSPT1 was predicted using TargetP2.0 (Armenteros et al. 2019), and the corresponding nucleotide sequence was removed to improve the expression efficiency. The TP-truncated coding sequence was codon optimized and synthesized by Eurofins Genomics (Tokyo, Japan). A yeast expression backbone was generated based on the pRS415 plasmid (Sikorski and Hieter 1989). The G418 resistance cassette (PGK1p–G418r–PGK1t) was amplified from pPGKNEO2 (Yoshida et al. 2007) and inserted into pRS415 using the In-Fusion HD cloning kit (Takara Bio), resulting in pRS415-G418r. Subsequently, the GAL1 promoter and TPI1 terminator regions were amplified from pYES2 (Thermo Fisher Scientific) and from a synthetic TPI1t fragment (obtained from the Saccharomyces Genome Database), respectively, and assembled into the pRS415-G418r backbone by In-Fusion cloning to generate pRS415-GAL1p-TPI1t-G418r, which served as the yeast expression vector for subsequent integration of GgBSPT1. The TP-truncated GgBSPT1 coding sequence was amplified using KOD One PCR Master Mix (Toyobo) and cloned into the linearized pRS415-GAL1p-TPI1t-G418r vector using the In-Fusion HD Cloning Kit (Takara Bio). Finally, the resulting construct was used for heterologous expression in *S. cerevisiae*.

### Transient expression and enzyme preparation from *N. benthamiana*

Transient expression and subsequent enzyme preparation in *N. benthamiana* were performed essentially as previously described (Karamat et al. 2014), with minor modifications. Expression plasmids carrying each PT gene and a plasmid encoding the silencing suppressor P19 were introduced into *Agrobacterium tumefaciens* strain LBA4404 by electroporation (Mattanovich et al. 1989). Transformed LBA4404 cultures expressing each PT and P19 were grown overnight at 28 °C, resuspended in sterile water, and combined before infiltration. Leaves of *N. benthamiana* were infiltrated on the abaxial surface, and infiltrated leaves were harvested 4 days later for enzyme preparation. Membrane fractions were isolated as follows. Approximately 4 g of infiltrated leaves were homogenized in 0.1 M potassium phosphate (KPi) buffer (pH 7.0) supplemented with 10 mM DTT, protease inhibitors, and PVPP (0.1 g per g fresh weight). The homogenate was centrifuged at 10,000 × g for 3 min at 4 °C, and the supernatant was filtered twice through Miracloth (Merck KGaA). The filtrate was then centrifuged at 20,000 × g for 1 h at 4 °C to obtain the microsomal fraction. The resulting pellet was resuspended in 100 mM Tris-HCl buffer (pH 7.5) containing 1 mM DTT and used as the crude membrane enzyme preparation for in vitro assays.

### Preparation of microsomal fractions from *S. cerevisiae*

Constructed GgBSPT1 expression vectors were introduced into *S. cerevisiae* BY4741 using the lithium acetate method (Gietz and Woods 2002). Transformants were precultured in YP medium containing 2 % glucose at 30 °C for 24 h and subsequently transferred to YP medium containing 2 % galactose for an additional 24 h to induce protein expression. Cells were disrupted with 0.5-mm glass beads using a Multi-Beads Shocker (Yasui Kikai). Microsomal fractions were prepared by centrifugation at 20,000 × g for 1 h at 4 °C, and the resulting pellet was resuspended in 100 mM CAPS buffer (pH 9.0) supplemented with 1 mM DTT and protease inhibitor. Protein concentrations were adjusted to 5 mg mL⁻¹.

### *In vitro* enzyme assays

Standard enzyme reactions were conducted in a total volume of 100 µL containing 0.2 mM isoliquiritigenin, 0.2 mM DMAPP, 10 mM MgCl₂, and 85 µL microsomal protein. Reaction mixtures were then incubated at 28 °C for 16 h for assays using *N. benthamiana* fractions, at 20 °C for 2 h for substrate specificity assays, and for 10 min for optimum-condition assays using *S. cerevisiae* fractions. For pH optimization, reactions were performed using three buffer systems: 0.1 M MES for pH 5.5–7.0, 0.1 M Tris-HCl for pH 7.0–9.0, and 0.1 M CAPS for pH 9.0–11.0. Crude membrane enzyme preparations were adjusted at pH 0.5 intervals within each range. Regarding temperature optimization, reactions were carried out at 5–40 °C in 5 °C increments. Reactions were terminated by addition of an equal volume of 2.8 N HCl and extracted with ethyl acetate. Finally, dried extracts were dissolved in methanol for LC–MS/MS analysis.

### HPLC and LC–MS/MS analysis

HPLC analyses were performed using a Shimadzu Prominence LC-20A system (Shimadzu, Kyoto, Japan) equipped with an XBridge C18 column (3.5 µm, 2.1 × 100 mm; Waters Corp., MA, USA), with the column oven maintained at 40 °C. Compounds were separated using a linear gradient of solvent B (0.1 % formic acid in acetonitrile) in solvent A (0.1 % formic acid in water) as follows: (min, %B) = (0, 10), (15, 95), (20, 95), (20.1, 10), and (27, 10), at a flow rate of 0.2 mL min⁻¹. For quantitative and qualitative analyses, mass spectrometric detection was performed using a Sciex 4500 QTrap mass spectrometer (AB Sciex LLC, MA, USA) with electrospray ionization in negative ion mode. For quantitative analysis, compounds were monitored in multiple reaction monitoring (MRM) mode. The ionspray voltage was set to −4.5 kV, the source temperature to 500 °C, curtain gas to 20 psi, ion source gas 1 to 50 psi, and ion source gas 2 to 30 psi. The declustering potential, entrance potential, collision energy, and collision cell exit potential were optimized for each analyte. MRM transitions were set for glabridin (m/z 323.1 > 201.3), isobavachalcone and isobavachin (m/z 323.0 > 119.0), and 4-hydroxylonchocarpin (m/z 321.0 > 119.0). For qualitative analysis, mass spectra were acquired in full-scan mode over an m/z range of 200–1000. The ionspray voltage was set to −4.5 kV, the source temperature to 450 °C, curtain gas to 30 psi, ion source gas 1 to 50 psi, and ion source gas 2 to 80 psi. Compound identities were further confirmed by comparison of precursor ion m/z values with those of authentic standards.

### Bioinformatic analysis of subcellular localization and transmembrane domains

Subcellular localization was predicted using DeepLoc-2.1 (Ødum et al. 2024), and transmembrane domains were predicted using DeepTMHMM 1.0 (Hallgren et al. 2022, bioRxiv).

### Statistical analysis

Data are presented as mean ± SD from three independent biological experiments. Data from YE and NC groups were analyzed using Welch’s one-way ANOVA followed by Games–Howell post hoc tests for multiple comparisons where appropriate. For comparisons between groups (YE vs. NC) at each time point, Welch’s t-test with Holm’s correction for multiple testing was applied. For enzyme activity assays, statistical significance was assessed by one-way ANOVA followed by Tukey’s multiple comparison test. Differences were considered statistically significant at p < 0.05.

## Results

### Accumulation of prenylated flavonoids by elicitation in cultured *G. glabra* roots

Cultured *G. glabra* roots were treated with YE, and time-course changes in prenylated flavonoid accumulation were monitored by LC–MS/MS analysis. As shown in Fig. 1, various prenylated flavonoids including chalcones, flavanones, and isoflavonoids were strongly induced. In particular, in the YE-treated group, isobavachin, 4-hydroxylonchocarpin, and glabridin exhibited clear time-dependent increases, reaching 3.4 ± 0.3, 7.6 ± 1.1, and 1.8 ± 0.2 μg g⁻¹ FW, respectively, on day 7 (Fig. 1b - d). In contrast, isobavachalcone (3′-prenylisoliquiritigenin) reached a high level (60 ± 9 μg g⁻¹ FW) as early as day 1 after treatment and remained essentially unchanged on days 3 and 7 (Fig. 1a). Throughout the experimental period, the accumulation levels of all four prenylated isoflavonoids were consistently higher in YE-treated roots than in NC. Only glabridin showed a transient decrease between day 0 and day 1, which may reflect stress responses associated with root excision. From the time-course profiles, induction of the three former compounds proceeded gradually, whereas isobavachalcone was induced rapidly, with strong accumulation occurring within 1 day after YE treatment. Notably, on day 7, isobavachalcone levels were approximately 9- to 37-fold higher than those of the other three prenylated flavonoid derivatives.

**Fig. 1.**
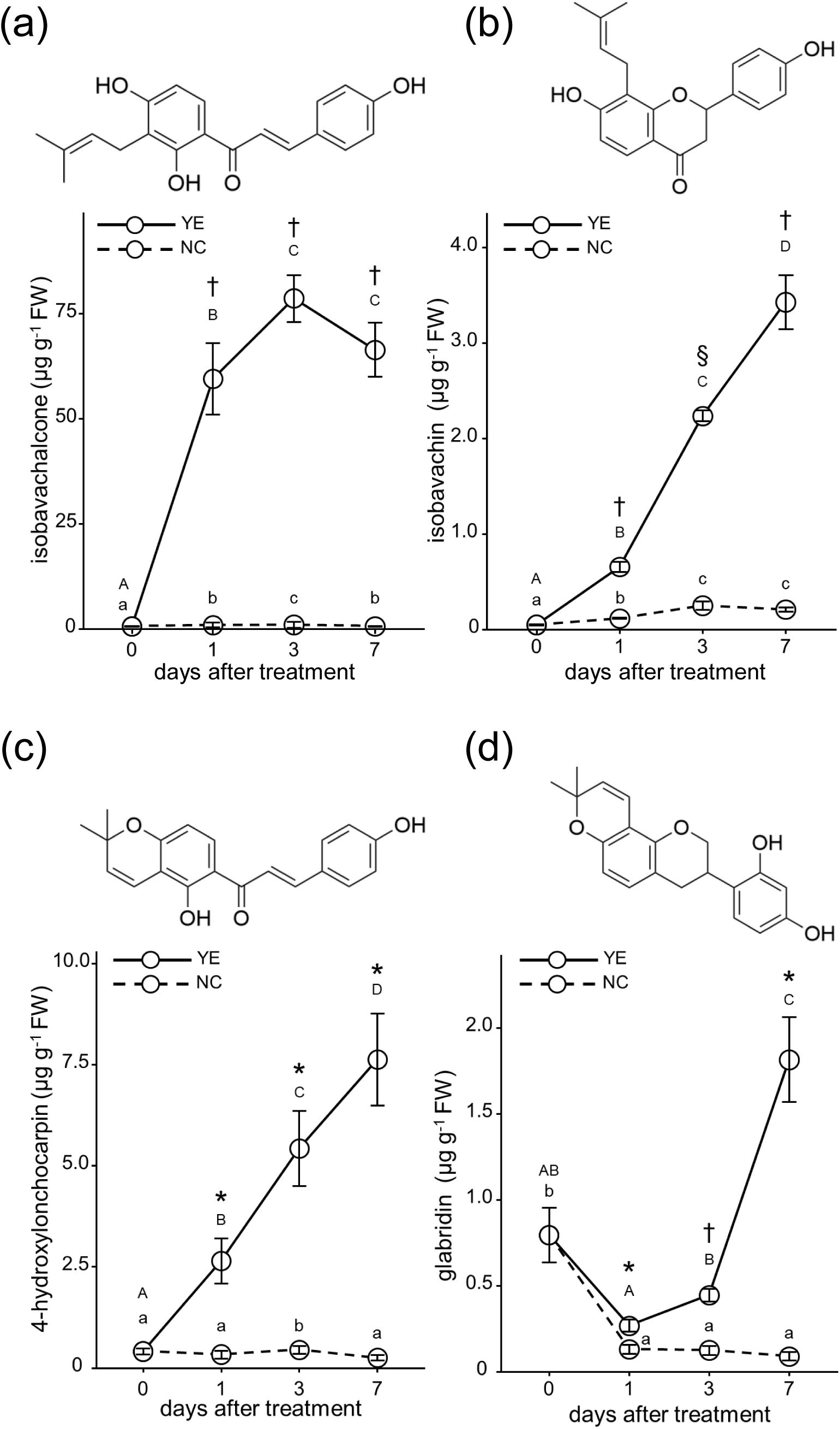
Time-course accumulation of prenylated and cyclized prenyl flavonoids in cultured *G. glabra* roots following stimulation Time-course changes in metabolite levels were monitored in cultured *G. glabra* roots treated with YE or NC, and concentrations were quantified by LC–MS/MS. Accumulation in the YE and NC groups is represented by solid and dashed lines, respectively. Chemical structures of the analyzed compounds are shown. Prenylated flavonoids: (a) isobavachalcone and (b) isobavachin. Cyclized prenyl flavonoids: (c) 4-hydroxylonchocarpin and (d) glabridin. Data are presented as mean ± SD (n = 3). Different letters indicate significant differences among time points within each treatment group (lowercase for NC and uppercase for YE). Symbols indicate significant differences between YE and NC at the same time point (*: p < 0.05; †: p < 0.01; §: p < 0.001). Statistical significance was assessed using Welch’s one-way ANOVA followed by Games–Howell post hoc tests for within-group comparisons and Welch’s t-test for between-group comparisons.

### Prenylation activity of crude enzyme extracts from cultured *G. glabra* roots

Because isobavachalcone showed the highest accumulation among the analyzed compounds, we next examined whether the corresponding prenylation reaction toward chalcone substrates occurs in cultured roots. Microsomal fractions were prepared from YE-elicited cultured *G. glabra* roots and used for enzyme assays with isoliquiritigenin as the acceptor substrate and DMAPP as the prenyl donor in the presence of Mg²⁺. Upon addition of isoliquiritigenin to the incubation mixture, a new peak (designated product 1) was detected by HPLC analysis (Fig. 2a). The retention time of product 1 matched that of authentic isobavachalcone. No corresponding peak was observed in the negative control, in which microsomal fractions were incubated without substrate (Fig. 2a). MS analysis further confirmed the identity of product 1 as isobavachalcone by demonstrating an identical m/z value to that of the authentic standard (Fig. 2b).

**Fig. 2.**
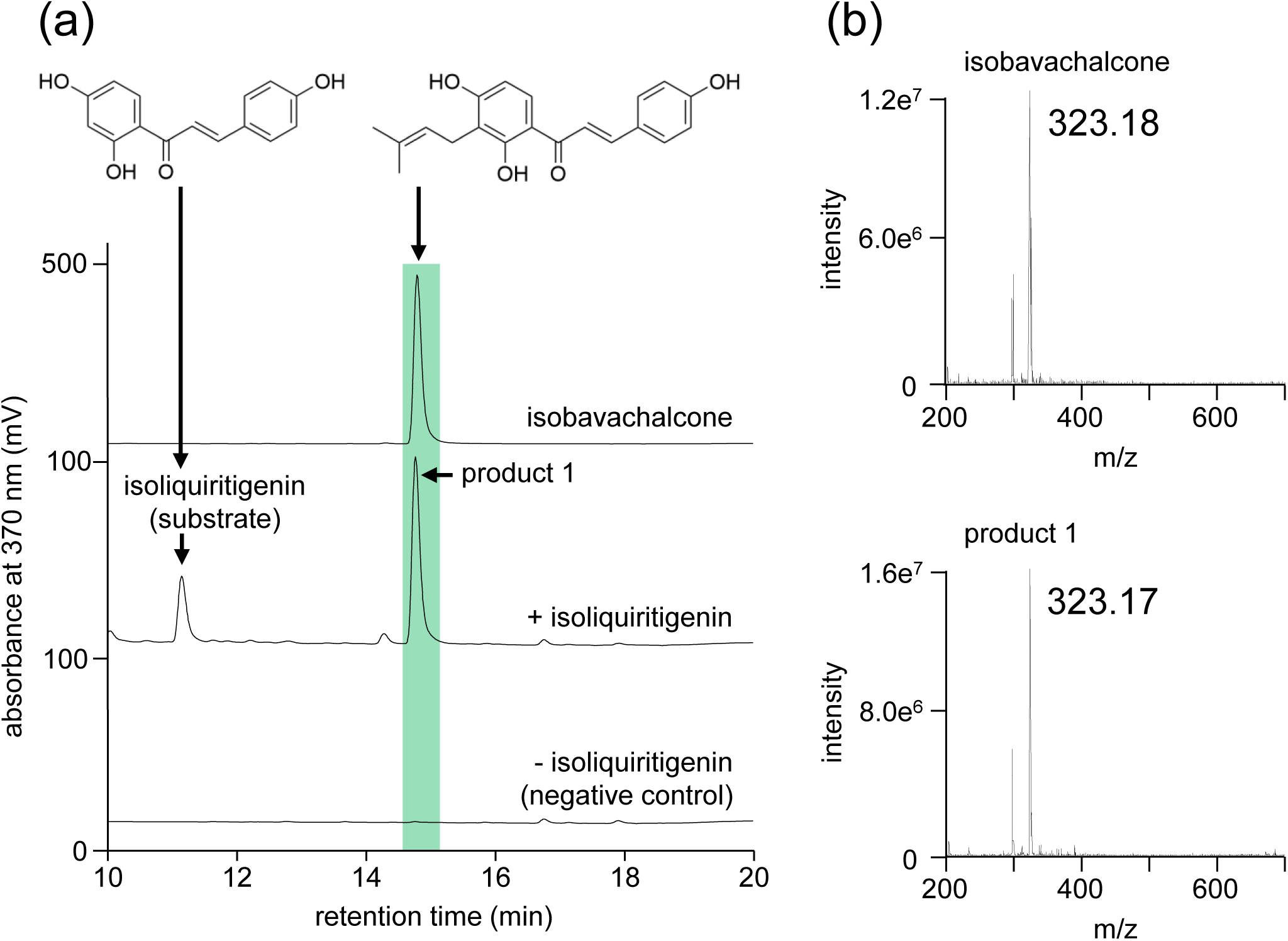
Prenylation activity of microsomal fractions from YE-elicited cultured *G. glabra* roots Microsomes prepared from YE-elicited roots were incubated with isoliquiritigenin as the acceptor substrate and DMAPP as the prenyl donor to evaluate prenylation activity. Reaction products were analyzed by (a) HPLC and (b) LC–MS. The full assay consisted of microsomal reactions supplemented with isoliquiritigenin (+ isoliquiritigenin), and the negative control consisted of microsomal reactions without substrate addition (– isoliquiritigenin).

### Cloning of PT cDNA in *G. glabra*

Because genomic and RNA-seq resources for *G. glabra* are limited, we first performed an in silico search for PT candidates using publicly available RNA-seq datasets of *G. uralensis*. Eight candidate PT genes were identified and tentatively designated GuPT1 to GuPT8. Full-length cDNA sequences of each candidate were synthesized and cloned into a plant expression vector. These constructs were transiently expressed in *N. benthamiana* via *Agrobacterium*-mediated infiltration, and microsomal fractions were subsequently prepared. Enzyme assays were conducted for each GuPT using isoliquiritigenin and DMAPP as prenyl acceptor and donor substrates, respectively, in the presence of Mg²⁺, with membrane fractions from *N. benthamiana* serving as crude enzyme preparations. Among the candidates, GuPT1 and GuPT3 catalyzed dimethylallyl transfer to isoliquiritigenin. In LC–MS analysis, a major reaction product (product 2) coeluted with authentic isobavachalcone and exhibited a molecular ion at m/z 323 identical to that of isobavachalcone, indicating that these enzymes function as isoliquiritigenin:DMAPP PTs (Fig. S1). In addition to this major product, GuPT1 generated two minor products (products 1 and 3 in Fig. S1), whereas GuPT3 produced one minor product (product 4 in Fig. S1); however, these minor products could not be identified.

As the synthesized PT of *G. urarensis* showed prenylation activity, we then tried to isolate homologous cDNAs from *G. glabra*. To isolate homologous cDNAs from *G. glabra*, a cDNA pool was synthesized from elicitor-treated cultured *G. glabra* roots, and PT genes were amplified from this template using primers designed based on GuPT1. A cDNA sharing 99 % nucleotide identity with GuPT1 was obtained and designated GgBSPT1 (Fig. 3). Figure 3 shows a phylogenetic tree constructed using representative UbiA-type PT proteins from Fabaceae, together with known PTs from Moraceae and Cannabaceae for comparison. UbiA PTs involved in secondary metabolism have evolved convergently and are therefore grouped according to plant families (Munakata et al. 2020). Within Fabaceae, PTs from *Glycyrrhiza* species constitute a closely related clade. Phylogenetic analysis indicated that GgBSPT1 (highlighted by black shading) is distinct from previously reported licorice PTs, namely GuA6DT (Li et al. 2014), GuILDT (Li et al. 2018), and the GgPT1 (Zhang et al. 2026), sharing 68 %, 78 %, and 70 % amino acid identities, respectively, which supports its novelty as being a new licorice PT. In silico analysis using DeepLoc-2.1 predicted the plastid localization of GgBSPT1 with high confidence (0.97). The N-terminal predicted transit peptide by TargetP 2.0 is indicated in green (Fig. S2). GgBSPT1 contains two conserved aspartate-rich motifs (N[Q/D]XXDXXXD and KD[I/L]XDX[E/D]GD), which are indicated in red (Fig. S2). These domains are characteristic of PT catalytic function, as well as nine predicted transmembrane α-helices that are indicated in blue (Fig. S2), consistent with typical UbiA-type PT features.

**Fig. 3.**
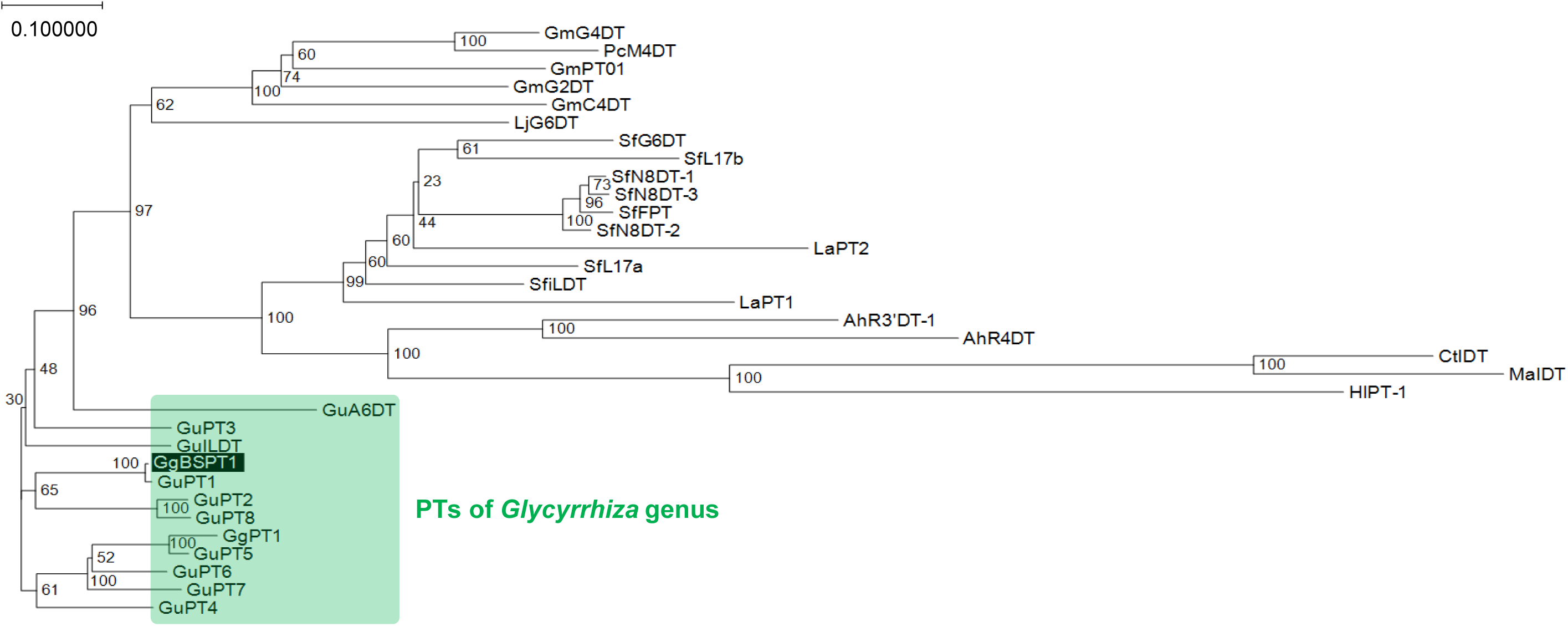
Phylogenetic analysis of GuPTs, GgBSPT1, and UbiA-type prenyltransferases from other plant species Protein sequences were aligned using ClustalW. A neighbor-joining phylogenetic tree was then constructed with GENETYX version 17. Bootstrap analysis was performed with 1,000 replicates, and bootstrap values are indicated at each node; branch lengths represent relative genetic distances. GgBSPT1 is indicated by black shading. The clade containing *Glycyrrhiza* PTs is highlighted in green. Abbreviations of protein sequences and corresponding accession numbers are as follows: SfN8DT-1 (AB325579), SfN8DT-2 (AB370330), SfL17a (AB371287), SfL17b (AB370329), GmG4DT (AB434690), SfN8DT-3 (AB604222), SfG6DT (AB604224), SfiLDT (AB604223), HlPT-1 (AB543053), LaPT1 (JN228254), SfFPT (KC513505), MaIDT (KM262659), CtIDT (KM262660), GuA6DT (KJ123716), GmC4DT (LC140927), GmG2DT (LC140930), GuILDT (KR139751), PcM4DT (MH626730), LjG6DT (KX228696), AhR4DT (KY565244), AhR3′DT-1 (KY565245), GmPT01 (NP_001335591), and LaPT2 (AWK21939).

### Enzymatic activity of GgBSPT1 expressed in *N. benthamiana*

To verify enzymatic function, full-length GgBSPT1 was transiently expressed in *N. benthamiana* by agroinfiltration. Membrane fractions prepared from the transient transformant leaves were subjected to enzyme assays using isoliquiritigenin as the acceptor substrate and DMAPP as the prenyl donor in the presence of Mg²⁺. Two reaction products (product 1 and 2) were detected. The major peak (product 2) corresponded precisely to authentic isobavachalcone in both retention time and m/z value (Fig. 4). These results demonstrate that GgBSPT1 catalyzes formation of isobavachalcone, which was a similar result as GuPT1. Although product 1 did not coelute with authentic isobavachalcone or bavachalcone standards, it displayed the same molecular ion (m/z 323) as isobavachalcone, indicating incorporation of a single prenyl group. This observation suggests that product 1 is a positional isomer of isobavachalcone that differs in the site of prenylation.

**Fig. 4.**
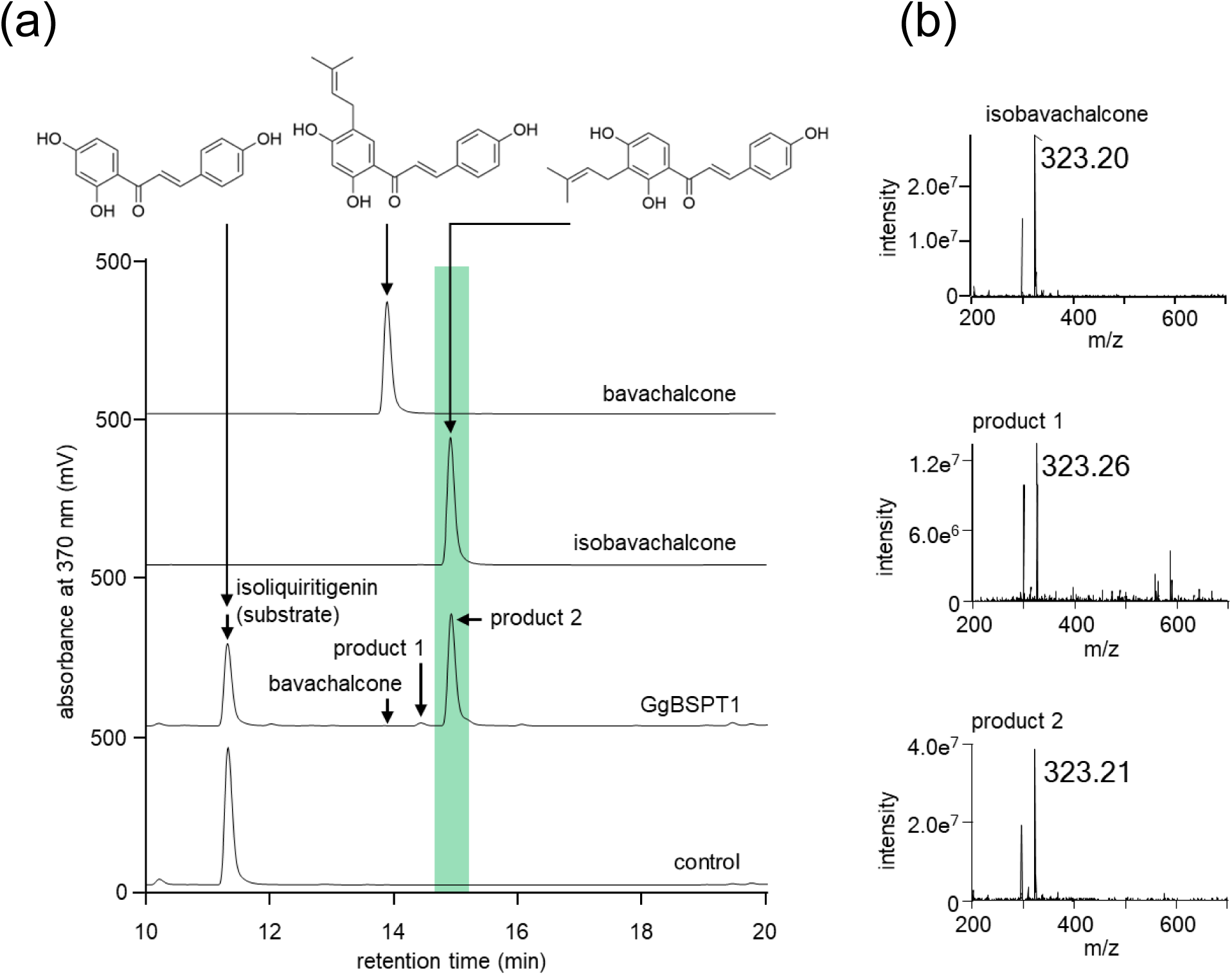
Prenylation activity toward isoliquiritigenin of microsomes prepared from *N. benthamiana* expressing GgBSPT1 Reactions were performed with isoliquiritigenin and DMAPP as substrates using microsomes from *N. benthamiana* expressing full-length GgBSPT1. The control represents the complete assay using microsomes from *N. benthamiana* expressing the empty vector. Reaction products were analyzed by (a) HPLC and (b) LC–MS. In LC–MS analysis, the expected deprotonated molecular ion ([M–H]⁻) of isobavachalcone was detected at m/z 323.

### Biochemical characterization of GgBSPT1 expressed in *S. cerevisiae*

For more detailed biochemical characterization, *S. cerevisiae* was selected as the heterologous host to facilitate higher-throughput analysis. A TP-truncated GgBSPT1 was expressed because TPs are generally unfavorable for yeast expression (Munakata et al. 2014). Enzyme assays using isoliquiritigenin and DMAPP in the presence of Mg²⁺ produced two clear reaction products (Fig. 5a). The major product was identified as isobavachalcone (product 2), confirming the functional expression of GgBSPT1 in yeast (Fig. 5a). As observed for GgBSPT1 expressed in *N. benthamiana*, the minor product (product 1) eluted earlier than the major product and could not be identified due to limited quantity and absence of an authentic standard.

**Fig. 5.**
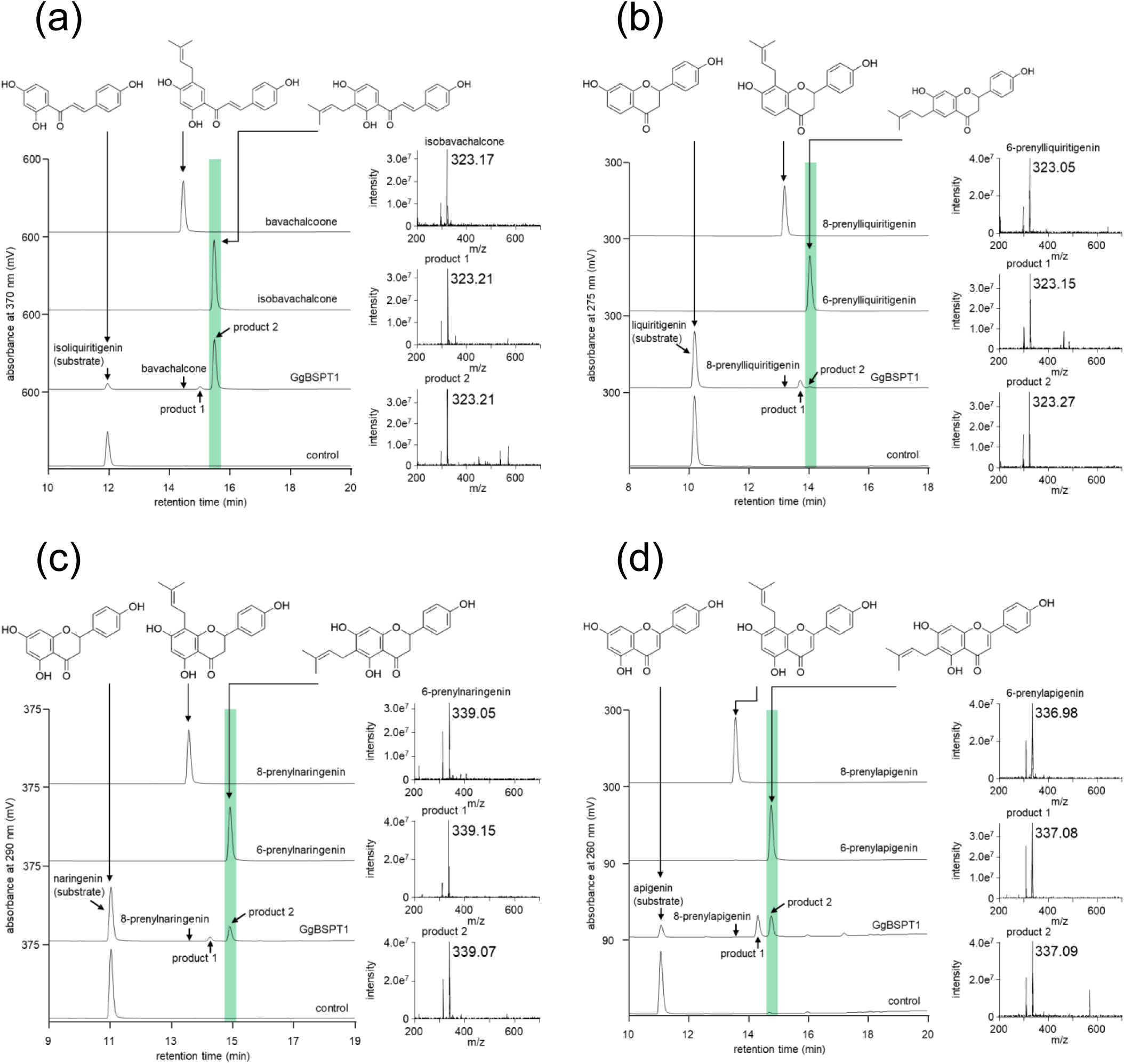
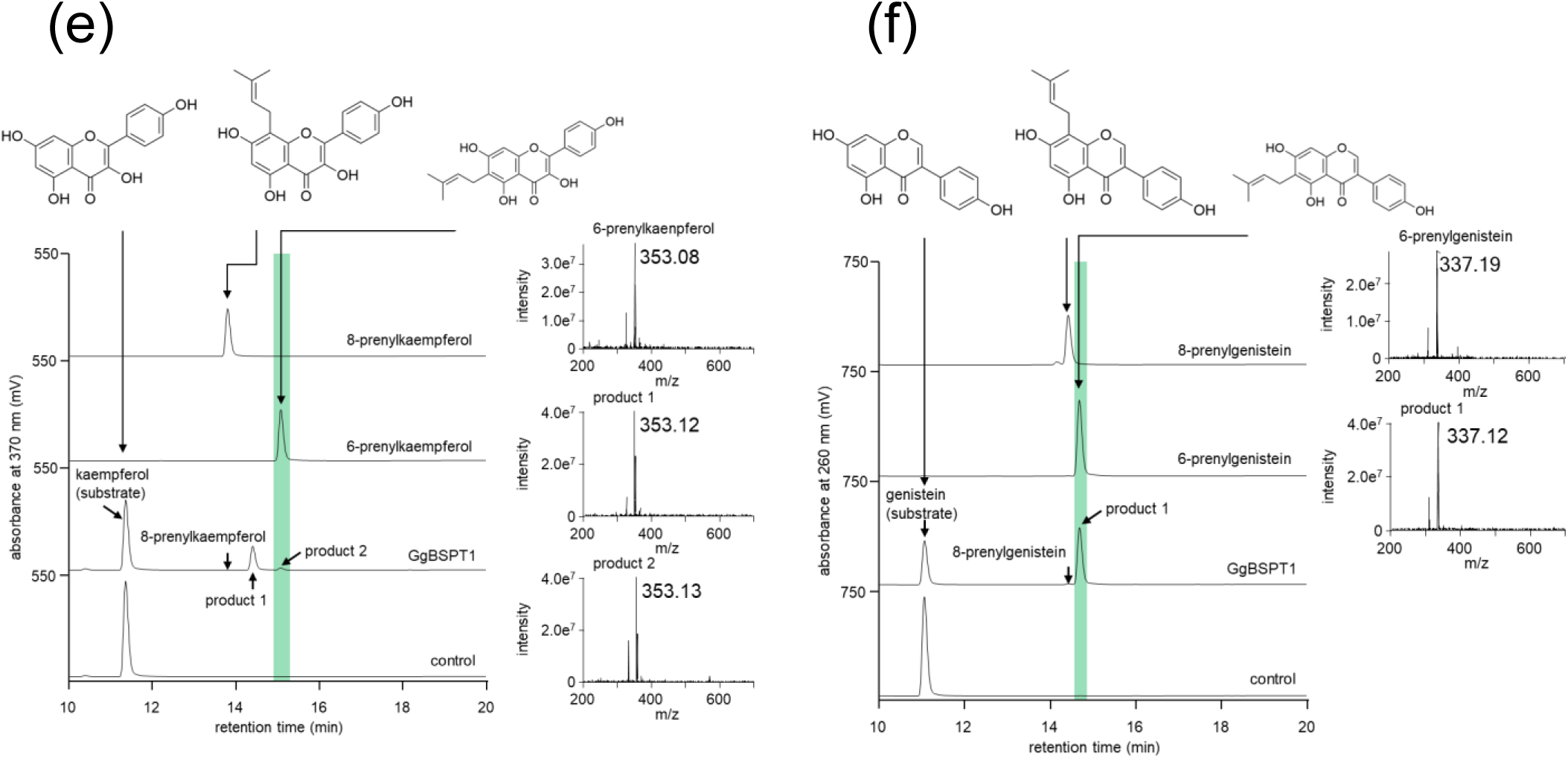
Substrate specificity of GgBSPT1 using microsomes prepared from transgenic *S. cerevisiae* Enzyme reactions were conducted with the following prenyl acceptor substrates: (a) isoliquiritigenin, (b) liquiritigenin, (c) naringenin, (d) apigenin, (e) kaempferol, and (f) genistein, in the presence of DMAPP using microsomes from *S. cerevisiae* expressing TP-truncated GgBSPT1. HPLC chromatograms and mass spectra of the reaction products are shown. Expected deprotonated molecular ions ([M–H]⁻) of each prenylated product were detected by MS. The control represents products obtained from the complete assay using microsomes expressing the empty vector.

Substrate specificity assays performed under optimal conditions (described below) revealed that GgBSPT1 possesses unusual catalytic properties among plant PTs. Most plant PTs exhibit strict substrate and product specificity, with precise control of substrate recognition and prenylation position. In contrast, GgBSPT1 catalyzed prenylation of chalcone, flavanone, flavone, flavonol, and isoflavone scaffolds (Fig. 5). For substrates with identifiable products (Fig. 5), GgBSPT1 consistently introduced the prenyl group at structurally equivalent positions across different flavonoid backbones. Isoliquiritigenin (chalcone) was prenylated at the 3′ position; liquiritigenin and naringenin (flavanones) were prenylated at the 6-position; apigenin (flavone) was prenylated at the 6-position; kaempferol (flavonol) yielded a major unidentified product together with a minor 6-prenylated product; and genistein (isoflavone) was predominantly prenylated at the 6-position. These reaction products matched authentic standards in retention time and m/z (Fig. 5). In addition, GgBSPT1 accepted 14 further flavonoid substrates, demonstrating broad acceptor specificity (Table 1; Fig. S4). Such extensive substrate promiscuity is highly unusual among plant PTs. Donor specificity assays indicated a strict preference for DMAPP, whereas GPP and FPP were not utilized. Further biochemical characterization showed that pH profiling using three buffer systems identified an optimum at pH 9.0–9.5 (Fig. S5a), and temperature profiling indicated an optimal range of 15–20 °C (Fig. S5b).

**Table 1.**
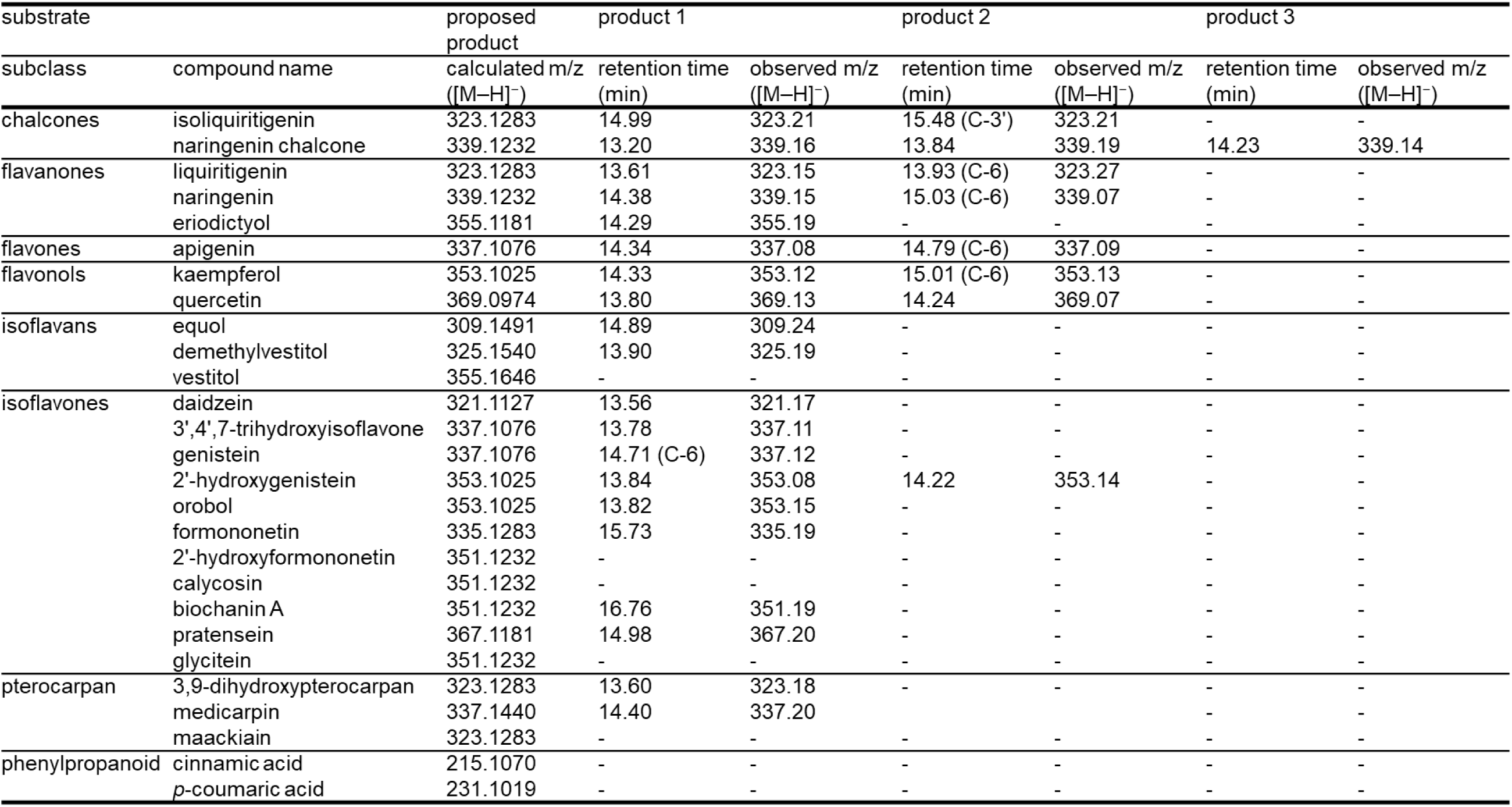
Summary of substrate and product specificity in the prenylation reaction catalyzed by GgBSPT1. The table summarizes substrates tested for GgBSPT1 activity using microsomes from *S. cerevisiae* expressing TP-truncated GgBSPT1, together with calculated m/z values of the expected deprotonated molecular ions ([M–H]⁻). Calculated m/z values are based on monoisotopic masses of deprotonated molecular ions ([M–H]⁻). For substrates exhibiting prenylation activity, retention time (min) and observed m/z ([M–H]⁻) values of each reaction product (products 1, 2, and 3) are listed. Prenylation position indicates the carbon atom (C-3′ or C-5′ of chalcones and C-6 or C-8 of flavonoids) assigned based on coelution with authentic standards and detection of the expected deprotonated molecular ions ([M–H]⁻) by LC–MS. For substrates showing no detectable prenylation activity, entries are indicated by “–.”

## Discussion

*Glycyrrhiza* species are widely used worldwide as sweeteners, food additives, and crude drugs. In particular, *G. glabra* and *G. uralensis* are the principal species consumed in the market. Major valuable compounds produced by *Glycyrrhiza* species include glycyrrhizin, a triterpene derivative, and diverse flavonoids, among which prenylated flavonoids are especially important because of their various biological activities (Yazaki et al. 2009). In contrast to extensive biosynthetic studies on glycyrrhizin (Chiyo et al. 2024), PTs catalyzing key prenylation steps in flavonoid biosynthesis have remained largely unidentified in *Glycyrrhiza* species. One possible reason is that candidate genes may be obscured by abundant nontarget transcripts highly expressed in RNA-seq datasets from intact plants. In the present study, we identified a novel PT (GgBSPT1) from *G. glabra* using elicitor-inducible adventitious root cultures. It has long been recognized that dedifferentiated cells, in vitro plants, and hairy roots can exhibit metabolite production profiles distinct from those of native plants (Hayashi et al. 1996; Kikowska et al. 2020; Kubica et al. 2022; Biswas et al. 2023). Consistent with these observations, elicited cultured *G. glabra* roots displayed pronounced alterations in prenylated flavonoid accumulation, with isobavachalcone showing the highest abundance among the analyzed compounds. The rapid induction of this compound suggests that chalcone-type prenylation represents an early step in the prenylated flavonoid pathway, responding more rapidly to elicitor signals than downstream modifications. Enhancement of PT activity following elicitor treatment has been reported in several plant species (Yoneyama et al. 2016; Sukumaran et al. 2018; Li et al. 2018), supporting the view that these enzymes contribute to stress-responsive metabolic plasticity. In agreement with these reports, YE-elicited cultured roots in our study showed markedly increased accumulation of prenylated flavonoids, indicating elevated prenylation activity under elicitation conditions. These observations prompted identification of the enzyme responsible for prenylated flavonoid formation.

Here, we characterized a previously unreported flavonoid PT, GgBSPT1, which differs from the GgPT1 identified in intact *G. glabra* (Zhang et al. 2026). Biochemical analysis revealed that GgBSPT1 exhibits unusually broad substrate scope and distinct regiospecific behavior (Fig. 5; Table 1; Fig. S4), thereby extending current understanding of PT diversity in plants (Fig. 5). Whereas plant PTs typically display high specificity for substrate recognition and prenylation position (Han et al. 2025), GgBSPT1 demonstrated a markedly broader catalytic profile. In general, PTs introduce prenyl groups at the ortho position relative to a phenolic hydroxyl group (Munakata et al. 2021). For six products whose prenylation sites were confirmed using authentic standards, prenylation occurred at the 3′ position in chalcones and at the 6-position in flavonoids. These positions are structurally equivalent, indicating that GgBSPT1 consistently targets the same aromatic carbon across diverse flavonoid scaffolds despite differences in backbone architecture. In isoliquiritigenin, naringenin, apigenin, kaempferol, and genistein, the prenylation occurred at a site flanked by two free hydroxyl groups, representing a favorable position for electrophilic substitution. In addition to the prenylation preference, GgBSPT1 exhibited broad substrate tolerance, catalyzing prenylation activity to 20 flavonoid-related structures across multiple subclasses (Table 1). Such extensive acceptor promiscuity exceeds that reported for previously characterized plant PTs and highlights GgBSPT1 as a metabolically versatile member of the UbiA superfamily. Collectively, these findings establish GgBSPT1 as an unusually flexible and catalytically active UbiA-type PT. GgBSPT1 connects elicitor-responsive metabolism with prenylated flavonoid biosynthesis in cultured *G. glabra* plants and likely contributes substantially to the chemical diversity observed in this species.

It has been reported that plant PTs involved in secondary metabolism have undergone taxon-specific evolution, such that phylogenetic relationships generally reflect plant taxonomy rather than enzymatic properties (Munakata et al. 2020). GgBSPT1 clusters within the Fabaceae PT clade, and the phylogenetic analysis (Fig. 3) further indicates that PTs identified in *G. uralensis* RNA-seq data, together with GgBSPT1 from *G. glabra*, constitute a *Glycyrrhiza*-specific clade with high amino acid sequence similarity. This pattern suggests the expansion of PT genes through multiple duplication events within this genus, likely driven by the central role of this enzyme family in diversification of prenylated flavonoids. A similar lineage-specific expansion of PT genes has been reported in *Lithospermum* species, as well as for other enzyme families such as acyltransferases in *Taxus* species (Kusano et al. 2019). Such duplication events may underlie the extensive chemical diversity of specialized metabolites in individual plant lineages.

From an industrial perspective, identification of novel enzymes derived from *Glycyrrhiza* species provides new opportunities for biotechnological production of valuable secondary metabolites. Recent work demonstrated that sequential introduction of biosynthetic enzymes into yeast can enable glabridin biosynthesis (Zhang et al. 2026). Nevertheless, the microbial production level of glabridin and related compounds are inefficient for the industrial production, but remains at the laboratory scale. Although these metabolites are currently obtained as plant extracts for use in cosmetics and nutraceuticals, de novo biosynthesis in engineered microorganisms has not yet reached commercially viable levels. This limitation arises from the complexity of the biosynthetic networks, limited enzyme activity or stability, insufficient precursor supply in vivo, and selection of appropriate microbial hosts (Han and Miao 2024). One promising strategy to address these challenges is identification of alternative metabolic branches, such as the prenylated isoflavonoid pathway revealed in elicited cultured *G. glabra* roots, followed by transfer of the corresponding genes into heterologous systems. Such an approach may expand the available repertoire of plant-derived biosynthetic genes and create new opportunities in synthetic biology of natural products. Taken together, these considerations emphasize the value of uncovering previously inaccessible enzymatic activities, and the usage of in vitro plant systems will provide a breakthrough to overcome these hurdles.

In summary, elicited cultured *G. glabra* roots enabled identification of a novel PT, GgBSPT1, that accepts diverse flavonoids as prenyl substrates and defines a previously uncharacterized branch of prenylated isoflavonoid metabolism. The broad substrate scope and flexible product specificity of GgBSPT1 provide new insight into metabolic versatility in *Glycyrrhiza* species. These findings further demonstrate the utility of in vitro plant systems for discovery of hidden biosynthetic functions and establish a basis for future reconstruction or engineering of prenylated phytochemical pathways in microbial or plant hosts.

## Declarations

### Author contribution statement

The research theme was conceived by A.K., who assumes responsibility for the integrity of the work. K.Y., K.O, and N.W. supervised the project and contributed to scientific discussions. A.K. performed enzymatic assays using *S. cerevisiae* and analyzed the data. O.K. constructed yeast expression vector. T.A. conducted metabolite accumulation analyses, cloned GgBSPT1, and performed enzymatic assays using *N. benthamiana*. Cultured *G. glabra* roots were established and maintained by N.Y., with technical assistance from K.M. J.H. identified candidate GuPT genes from transcriptome datasets, and K.N. contributed to sequence analysis. R.M. provided guidance on experimental design and enzymatic analyses. The manuscript was written by A.K., R.M., K.Y., and K.O. All authors have read and approved the final manuscript.

### Data availability

All data supporting the findings of this study are available within the manuscript and its supplementary materials. The nucleotide sequences of GuPT1-GuPT8 and GgBSPT1 have been deposited in the DDBJ database under accession numbers YABG01000001-YABG01000008 and LC925036, respectively.

## Acknowledgements

We thank Dr. Mareshige Kojoma (Health Sciences University of Hokkaido) for providing seeds of *G. glabra*.

