## supplementaly information for "An aromatic substrate prenyltransferase involved in the chemical diversification of flavonoids in *Glycyrrhiza glabra*"

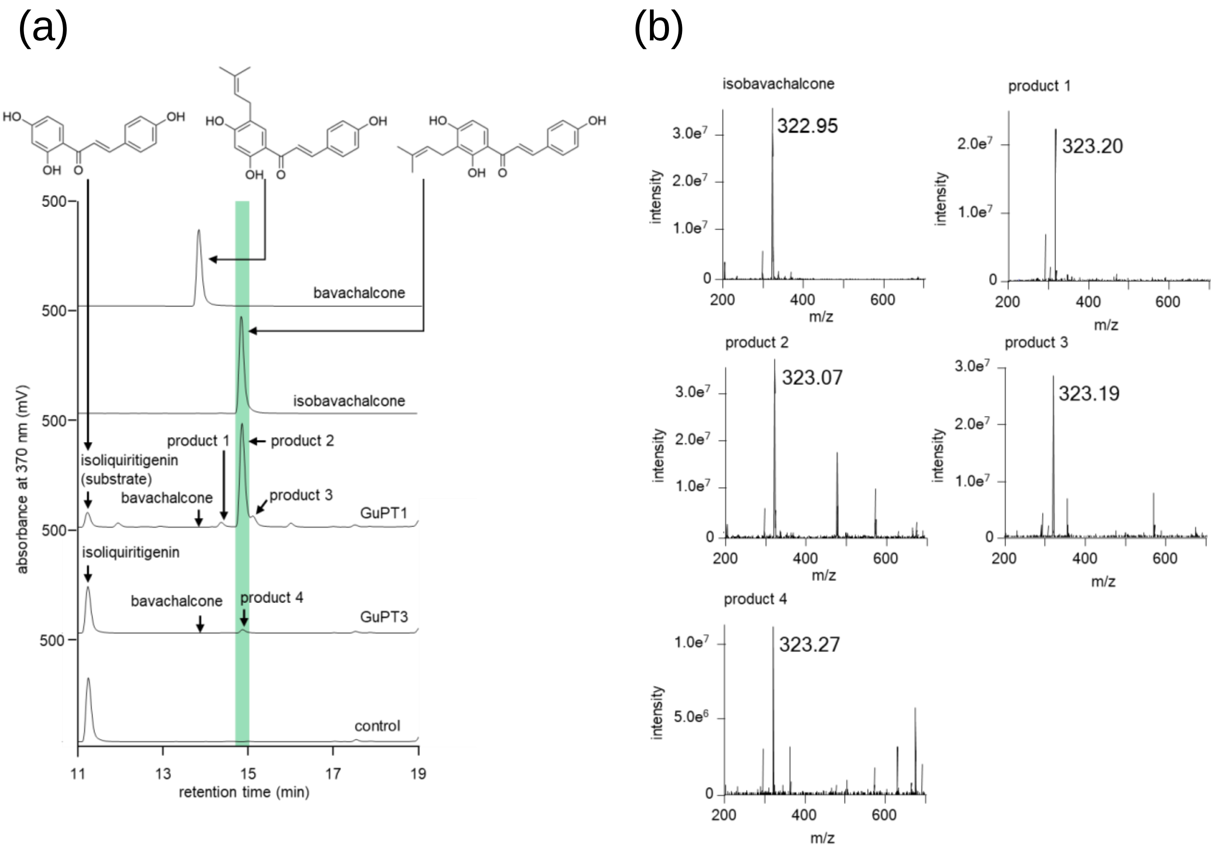


**Fig. S1** Prenylation activity toward isoliquiritigenin of microsomes prepared from *N. benthamiana* expressing GuPT1 or GuPT3

Reactions were performed with isoliquiritigenin and DMAPP using microsomes from *N. benthamiana* expressing GuPT1 or GuPT3. The control represents the complete assay using microsomes from *N. benthamiana* expressing the empty vector. Reaction products were analyzed by (a) HPLC and (b) LC–MS. In LC–MS analysis, the expected deprotonated molecular ion ($[M–H]^-$) of isobavachalcone was detected at m/z 323.

Putative transit peptide

MDSLIVGSFP KASSINSGGN LLRGENRTKS YYATSSYTPK ASLRKRKTQK EYNFLRSQQT

SLKHLYKGVE GGFTYQEYNR KYVVKTAPKT SFVSDPRSIE WKNILESVKT FLDAFYMFIT

Transmembrane domain — 1st D-rich motif

PYSVIATVLS IISACLLAVE KLSDISPLFF TGVLQAIIPH LFMSIYVNGI NQLGDIEIDK

INKPYLPLAS GKISFTTGAI IVASSLILSL WLAWIVGSWP SIWALISIAM IWGAYSVNVP

LLRWKRYPVL AAMVIVGSFS IVFPIGYFLH MQTFVFKRPA FFSRPLIFAT TFTSFFSLVI

2nd D-rich motif

ALFKDIPDIE GDQAFGVQSF AARLGQKRVF WICISLLEMA YGVALLMGLT SSCLWSKTVT

VLGHTVLASV VLYRAKSVDL RSKASIVSFY MLIWKLLSVE YFLMPLVR 408 a.a.

**Fig. S2** Amino acid sequence and structural features of GgBSPT1
The full-length protein sequence is shown. The putative transit peptide, conserved aspartate-rich motifs, and transmembrane α-helices are indicated in green, red, and blue, respectively.

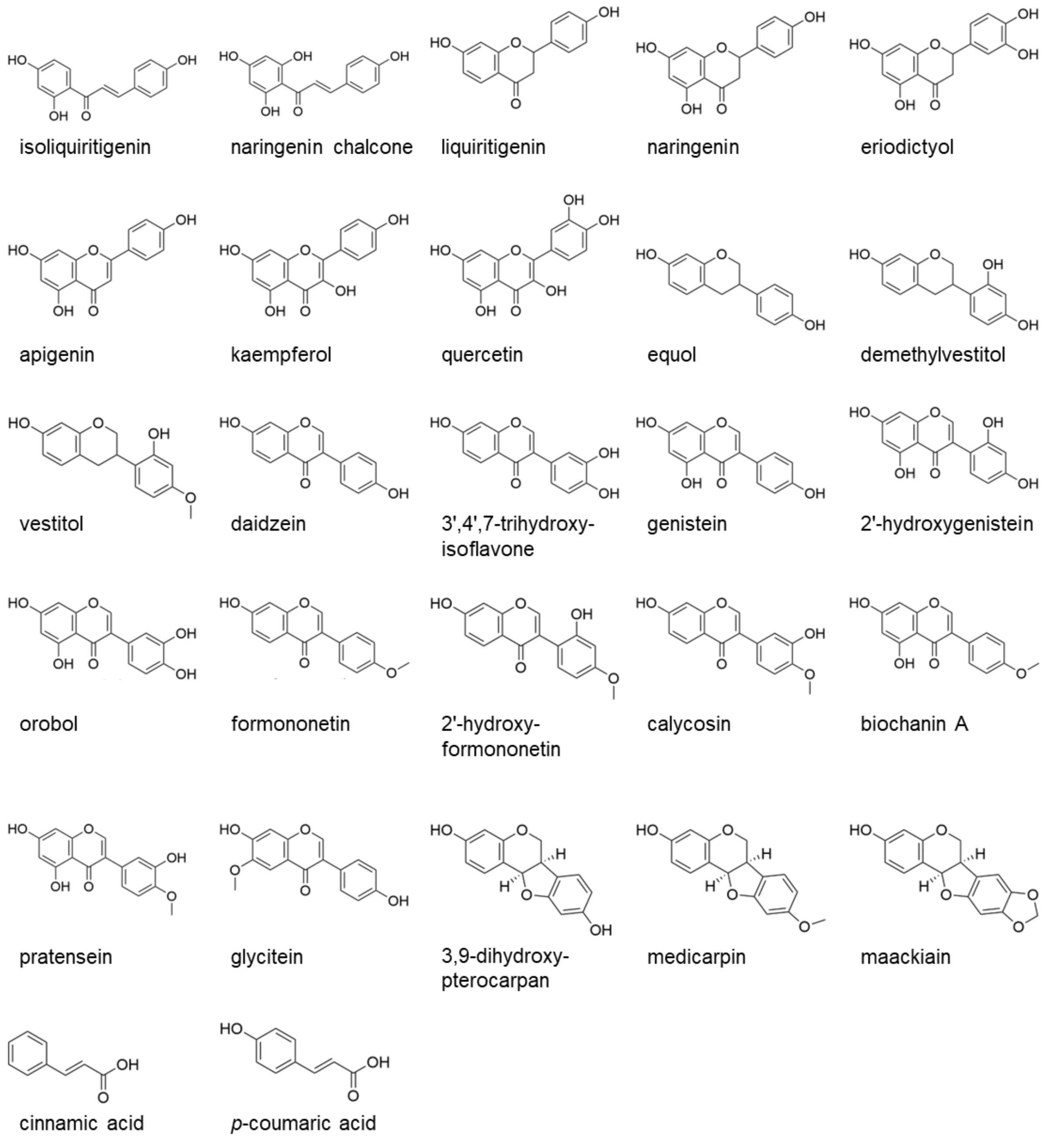


**Fig. S3** Chemical structures of flavonoid substrates used in GgBSPT1 enzyme assays
Names of the substrates are indicated in the figure. The depicted compounds correspond to those listed in Table 1.

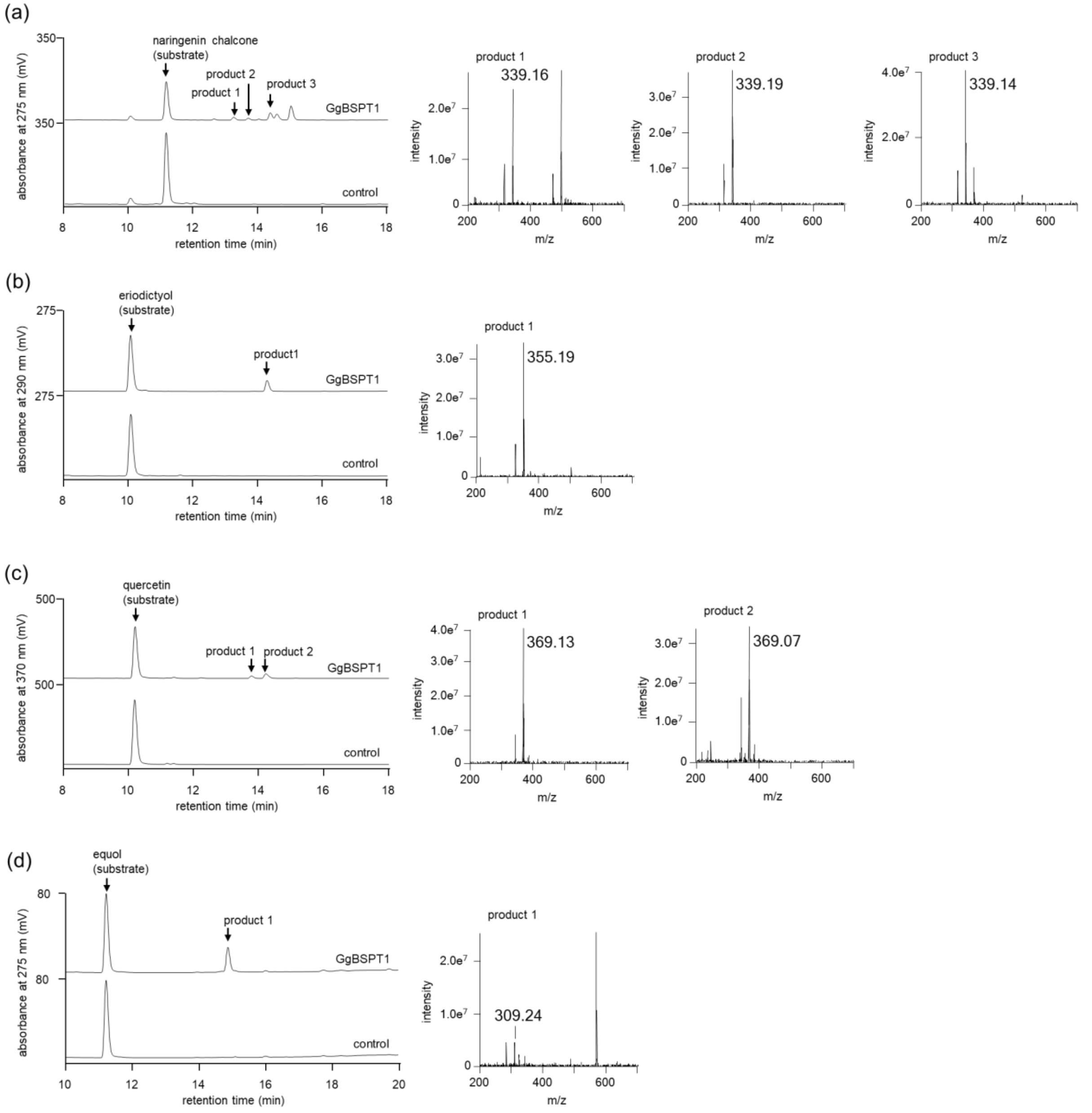


**Fig. S4** Extended GgBSPT1 substrate specificity using microsomes of transgenic *S. cerevisiae*
Reactions were performed with (a) naringenin chalcone, (b) eriodictyol, (c) quercetin, (d) equol, (e) desmethylvestitol, (f) daidzein, (g) 3',4',7-trihydroxyisoflavone, (h) 2'-hydroxygenistein, (i) orobol, (j) formononetin, (k) biochanin A, (l) pratensein, (m) 3,9-dihydroxypterocarpan, and (n) medicarpin in the presence of DMAPP, using microsomes from *S. cerevisiae* expressing TP-truncated GgBSPT1. HPLC chromatograms and mass spectra of enzymatic reaction products are shown. The control represents products obtained from microsomes expressing the empty vector. For products lacking authentic standards, formation was evaluated based on LC–MS detection of the expected deprotonated molecular ions ($[M–H]^–$).

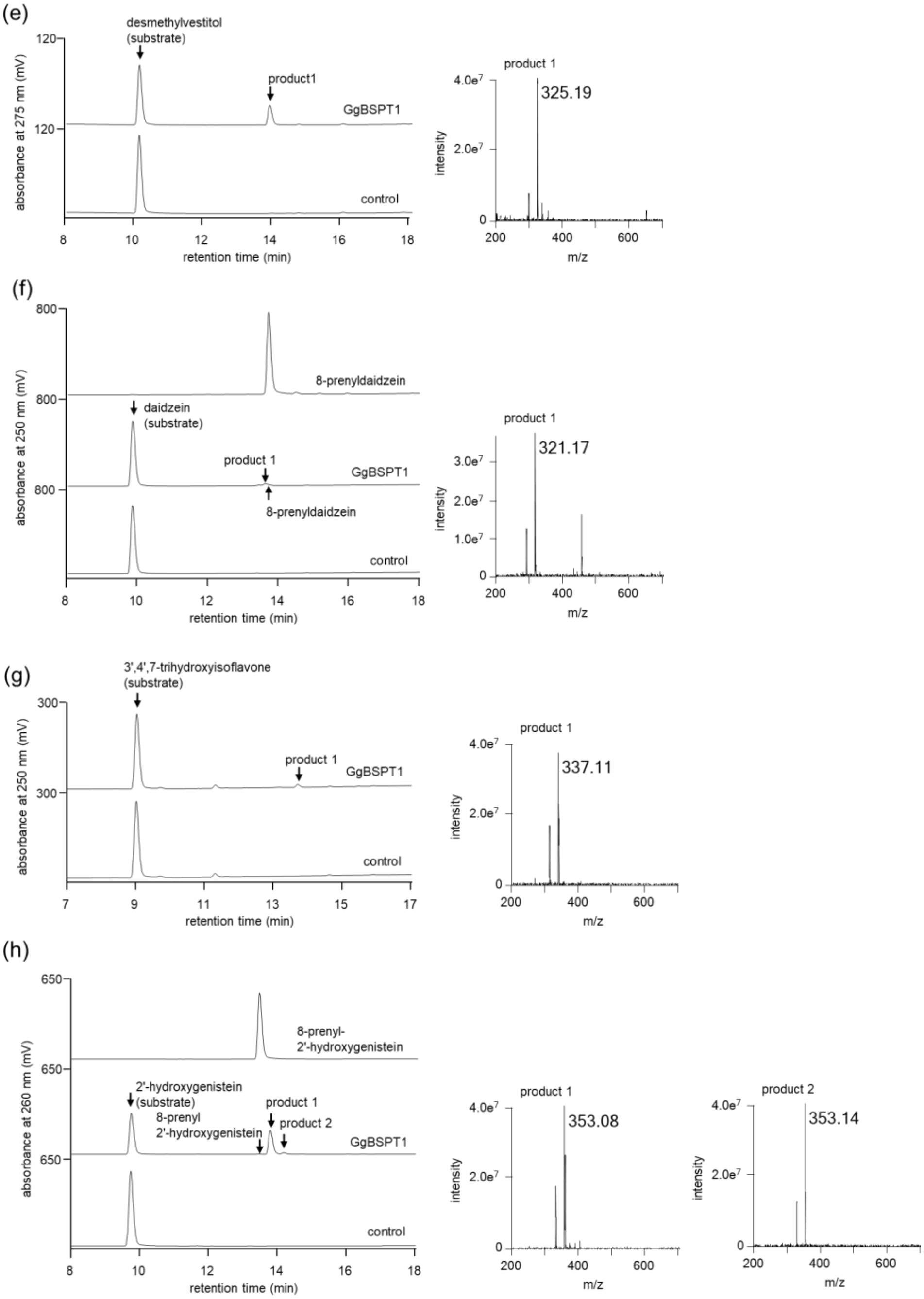
(e)
desmethylvestitol (substrate)
product1
GgBSPT1
control
absorbance at 275 nm (mV)
120
retention time (min)
product 1
325.19
intensity
4.0e7
2.0e7
m/z
(f)
800
8-prenyldaidzein
daidzein (substrate)
product 1
GgBSPT1
8-prenyldaidzein
control
absorbance at 250 nm (mV)
retention time (min)
product 1
321.17
3.0e7
2.0e7
1.0e7
intensity
m/z
(g)
3',4',7-trihydroxyisoflavone (substrate)
300
product 1
GgBSPT1
control
absorbance at 250 nm (mV)
retention time (min)
product 1
337.11
4.0e7
2.0e7
intensity
m/z
(h)
650
8-prenyl-
2'-hydroxygenistein
2'-hydroxygenistein (substrate)
8-prenyl
2'-hydroxygenistein
product 1
product 2
GgBSPT1
control
absorbance at 260 nm (mV)
retention time (min)
product 1
353.08
product 2
353.14
4.0e7
2.0e7
intensity
m/z

(i)

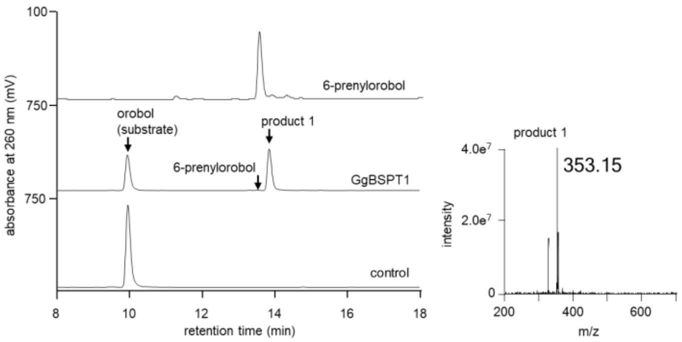
100
750
750
absorbance at 260 nm (mV)
6-prenylorobol
orobol
(substrate)
product 1
6-prenylorobol
GgBSPT1
control
8
10
12
14
16
18
retention time (min)
product 1
4.0e7
2.0e7
0
353.15
intensity
200
400
600
m/z

(j)

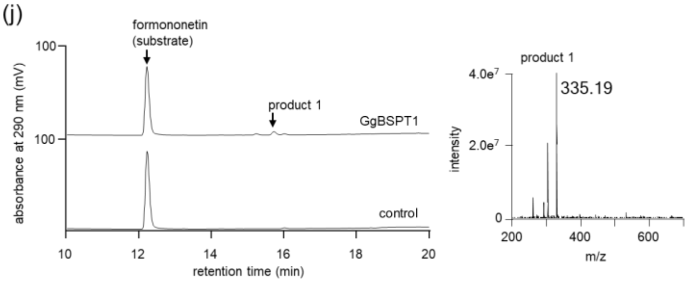
100
100
absorbance at 290 nm (mV)
formononetin
(substrate)
product 1
GgBSPT1
control
10
12
14
16
18
20
retention time (min)
product 1
4.0e7
2.0e7
0
335.19
intensity
200
400
600
m/z

(k)

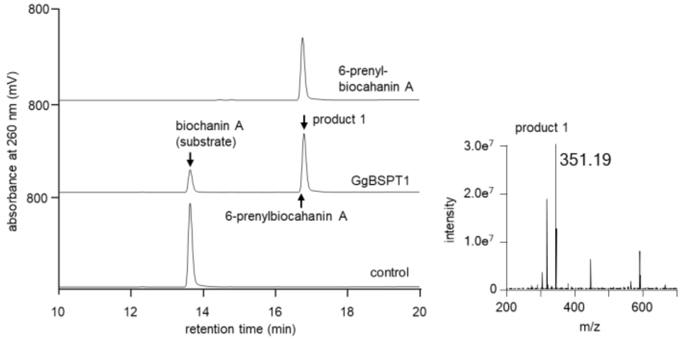
800
800
800
absorbance at 260 nm (mV)
6-prenyl-
biocahanin A
biochanin A
(substrate)
product 1
GgBSPT1
6-prenylbiocahanin A
control
10
12
14
16
18
20
retention time (min)
product 1
3.0e7
2.0e7
1.0e7
0
351.19
intensity
200
400
600
m/z

(l)

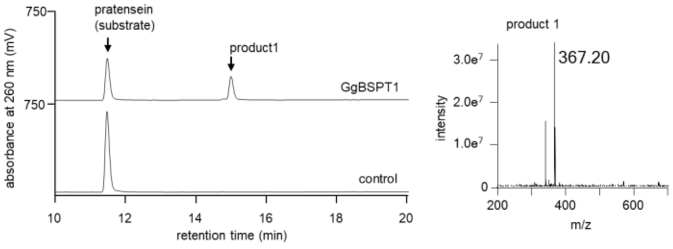
750
750
absorbance at 260 nm (mV)
pratensein
(substrate)
product1
GgBSPT1
control
10
12
14
16
18
20
retention time (min)
product 1
3.0e7
2.0e7
1.0e7
0
367.20
intensity
200
400
600
m/z

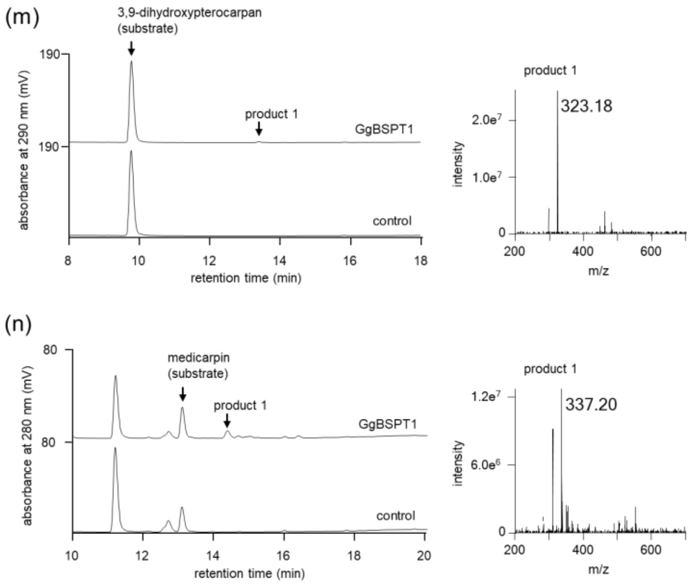
(m)
3,9-dihydroxypterocarpan
(substrate)
190
190
product 1
GgBSPT1
control
absorbance at 290 nm (mV)
8
10
12
14
16
18
retention time (min)
product 1
323.18
2.0e7
1.0e7
0
intensity
200
400
600
m/z
(n)
80
80
medicarpin
(substrate)
product 1
GgBSPT1
control
absorbance at 280 nm (mV)
10
12
14
16
18
20
retention time (min)
product 1
337.20
1.2e7
6.0e6
0
intensity
200
400
600
m/z

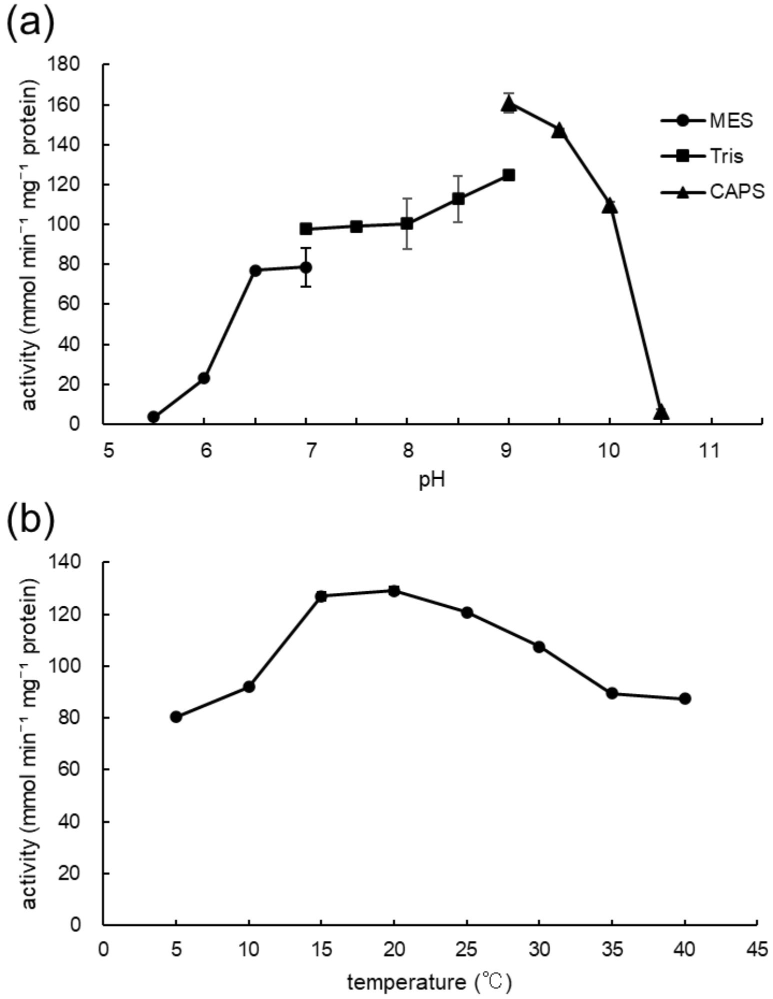


**Fig. S5** Optimal reaction conditions of GgBSPT1

Microsomes were prepared from *S. cerevisiae* expressing TP-truncated GgBSPT1. (a) Enzyme activity was measured across a pH range of 5.5–11. (b) Enzyme activity was measured at temperatures from 5℃ to 40 ℃ . Data are presented as mean ± SD (n = 3). MES, Tris, and CAPS buffers are indicated by circles, squares, and triangles, respectively.
